# Cohesin collisions maintain ordered nucleosome architecture at boundaries and promoters

**DOI:** 10.64898/2026.05.22.727261

**Authors:** Ramya Raviram, Guimei Jiang, Tom Schippke, Giulia Cova, Jane A Skok

## Abstract

Cohesin is best known for its role in loop extrusion, while nucleosome phasing at regulatory elements is usually attributed to local DNA-bound factors and remodelers. Here we identify a previously unrecognized role for cohesin-mediated extrusion in maintaining local nucleosome architecture at CTCF sites and transcription start sites. Using single-molecule nano-NOMe-seq during SCC1 depletion, cell-cycle progression and Sororin perturbation, we show that CTCF-bound sites contain distinct nucleosome architectures ranging from ordered CTCF-footprinted arrays to footprint-free nucleosomal and inaccessible configurations.. In unperturbed cells, ordered CTCF-footprinted nucleosome arrays were strongest at a boundary-enriched class of CTCF sites without regulatory elements. By contrast, CTCF sites overlapping regulatory elements showed stronger aggregate CTCF ChIP-seq signal despite weaker footprinting and less regular nucleosome phasing, indicating that boundary-like nucleosome architecture is not predicted by CTCF occupancy alone. At TSSs, promoter-proximal CTCF defined a distinct state balance: CTCF-positive promoters were enriched for accessible and footprinted configurations, whereas CTCF-negative promoters showed proportionately fewer footprinted states and were dominated by footprint-free phased arrays. Acute SCC1 depletion disrupted nucleosome organization at CTCF sites without regulatory elements and at promoters with promoter-proximal CTCF, despite retention of aggregate CTCF ChIP-seq signal at CTCF-bound sites. SCC1 depletion also altered nucleosome organization at promoters lacking promoter-proximal CTCF, highlighting that cohesin-dependent nucleosome patterning is not simply a CTCF-barrier phenomenon. Cell-cycle and Sororin analyses further separated extrusion-associated states from Sororin-stabilized post-replicative cohesin, highlighting that nucleosome order depends on effective cohesin-barrier encounters rather than cohesin occupancy alone. Together, these findings establish cohesin collisions as an active local mechanism that patterns nucleosomes at boundaries and promoters.

## Introduction

Mammalian genomes are folded by cohesin and CTCF into loops, contact domains, and insulating boundaries, yet the molecular state that converts this architecture into local chromatin organization and boundary function remains incompletely understood. Cohesin can extrude DNA loops, and in cells this activity is thought to continue until cohesin encounters boundary elements, many occupied by convergently oriented CTCF, thereby generating focal loops and topologically associating domains ^1–3^. Cohesin also accumulates at active promoters and regulatory elements, where it may encounter transcription factors, promoter-bound complexes, or other chromatin features that can impede or position extrusion independently of canonical CTCF boundaries. Acute depletion experiments have established that CTCF is required for CTCF-anchored loops and for local domain insulation, while cohesin depletion disrupts loop domains more broadly ^4^. However, disruption of loops and insulation does not always produce equivalent changes in steady-state transcription, indicating that contact maps, average CTCF occupancy, and aggregate accessibility do not fully capture the chromatin features relevant to boundary function. We therefore asked whether local nucleosome organization provides an additional readout of cohesin–CTCF boundary state ^5^.

Another key unresolved problem is why only a subset of CTCF-bound sites behave as strong insulating boundaries and whether the same local chromatin-ordering principles apply at regulatory elements that recruit or position cohesin in different ways. CTCF is not a uniform architectural signal: it has been implicated in insulation, transcriptional activation and repression, enhancer–promoter organization, imprinting, and other regulatory functions ^6^. Thus, CTCF binding must be interpreted within its local chromatin environment. Some CTCF sites lie within accessible, TF-rich promoter or enhancer regions, where CTCF occupancy is superimposed on transcription-factor binding, active chromatin, and promoter-associated accessibility. Others function as dedicated architectural barriers, where CTCF–cohesin encounters may be the dominant organizing event. Similarly, many TSSs are cohesin-associated even in the absence of promoter-proximal CTCF, suggesting that promoter chromatin organization may depend on cohesin positioning through CTCF-dependent and CTCF-independent mechanisms. In vitro reconstitution has further shown that CTCF is not simply a passive roadblock to loop extrusion, but an active, DNA-tension-dependent regulator of cohesin movement, direction switching, and loop shrinkage ^7^. These observations indicate that local chromatin organization depends not only on whether CTCF is bound, but on the type of chromatin barrier or regulatory context in which cohesin is positioned.

Local nucleosome organization provides a direct molecular readout of this context. CTCF sites are embedded within phased nucleosome arrays, and CTCF itself can help organize this local nucleosome architecture ^8^. However, nucleosome organization is not equivalent to CTCF occupancy. Recent single-molecule and genomic analyses indicate that motif grammar and transcriptional context can decouple CTCF binding from nucleosome phasing, and that nucleosome phasing regularity predicts insulation strength more reliably than CTCF ChIP intensity ^9^. This suggests a model in which a strong boundary is not simply a CTCF-bound site, but a CTCF-centered chromatin state: a configuration with a coherent CTCF footprint, phased flanking nucleosomes, and local cohesin engagement. These observations raise the possibility that boundary CTCF sites, enhancer/TSS-associated CTCF sites, and CTCF-independent TSSs occupy distinct single-molecule chromatin states, but whether cohesin helps maintain these states independently of CTCF occupancy remains unclear.

This distinction is difficult to resolve with CTCF depletion alone. CTCF sites are classically associated with phased flanking nucleosomes, but CTCF depletion removes more than the DNA-bound factor: it also removes the major barrier that positions cohesin. Consequently, loss of local order after CTCF depletion cannot distinguish direct CTCF-dependent nucleosome organization from loss of cohesin barrier encounters. This ambiguity motivated us to ask whether local nucleosome order at architectural boundaries and promoter-proximal regulatory elements depends on how cohesin is positioned, and whether acute depletion of SCC1/cohesin shifts individual chromatin fibers from ordered to disordered states. We therefore stratified CTCF sites based on their overlaps with regulatory elements and TSSs by promoter-proximal CTCF occupancy. This framework allowed us to test whether cohesin-dependent nucleosome ordering is restricted to architectural boundaries or extends across distinct cohesin - associated regulatory contexts.

Cohesin supports both loop extrusion and cohesion, but these activities are temporally and mechanistically separable. Its residence time and chromatin engagement change as cells progress from G1 into S/G2, when replication-coupled chromatin maturation and establishment of a more stable cohesin pool create distinct post-replicative chromatin states. Comparing boundary and promoter chromatin states across the cell cycle can therefore help distinguish organization linked to ongoing extrusion from that associated with Sororin-stabilized cohesive cohesin. Sororin is recruited to replication-acetylated cohesin and stabilizes cohesin on chromatin by antagonizing WAPL-mediated release ^10^. Sororin depletion in G2 can therefore test whether ordered nucleosome states at boundaries and promoters depend mainly on dynamic cohesin–barrier encounters or on Sororin-stabilized post-replicative cohesin.

Distinguishing these possibilities requires resolving chromatin organization on individual DNA molecules rather than relying on population-averaged measurements, an approach enabled by long read single-molecule methyltransferase-footprinting methods that capture nucleosome protection and accessibility on individual chromatin fibers ^11,12^. Population-averaged profiles can reveal an aggregate CTCF footprint or phased nucleosome pattern, but they cannot distinguish uniform CTCF binding in one chromatin environment from a mixture of molecules bound within distinct ordered, disordered, or accessible chromatin states. Nano-NOMe-seq combines nanopore long-read sequencing with exogenous GpC methyltransferase labeling, revealing chromatin accessibility and nucleosome protection on individual DNA molecules ^13^. We used this single-molecule information to define chromatin states separately across CTCF sites at or away from regulatory elements, and TSSs with or without promoter-proximal CTCF.

By pooling molecules across conditions before clustering, we quantified how acute SCC1/cohesin depletion, cell-cycle state, and Sororin perturbation redistributed single molecules among shared nucleosome-architecture classes within each regulatory context. This analysis showed that CTCF-bound sites and promoters are not organized as uniform chromatin classes, but instead occupy distinct configurations defined by CTCF footprint strength, nucleosome phasing, accessibility, and local regulatory context. In unperturbed cells, CTCF sites without regulatory elements displayed the sharpest CTCF footprints and most regular flanking nucleosome phasing, whereas CTCF-positive and CTCF-negative TSSs adopted distinct promoter-associated nucleosome architectures, including configurations with a positioned +1 nucleosome that may reflect paused or early elongation-associated promoter organization. Importantly, boundary-like nucleosome architecture was not predicted by aggregate CTCF ChIP-seq strength. Indeed, CTCF sites overlapping regulatory elements showed stronger CTCF signals despite weaker footprinting and less regular nucleosome phasing.

Acute SCC1 depletion disrupted nucleosome organization at CTCF sites without regulatory elements and promoters with proximal CTCF, despite retention of aggregate CTCF ChIP-seq signal. This was also observed at promoters lacking proximal CTCF. Thus, cohesin-dependent nucleosome patterning is not simply a CTCF-barrier phenomenon, but extends to promoter contexts in which cohesin may be positioned by additional regulatory features. Cell-cycle and Sororin perturbation analyses further distinguished Sororin-stabilized cohesive cohesin from dynamic, extrusion-competent cohesin, linking local nucleosome order most closely to effective cohesin encounters with CTCF sites, promoters, or other chromatin impediments. Together, these findings support a model in which cohesin collisions help maintain ordered nucleosome architecture at boundaries and promoters. In this view, local genome organization is defined not only by factor occupancy, but by the distribution of single-molecule nucleosome architectures within each regulatory context.

## Results

### Nano-NOMe-seq defines shared single-molecule chromatin states across conditions

In nano-NOMe-seq, intact nuclei are treated with GpC methyltransferase to label accessible DNA, followed by long-read nanopore sequencing to read rmethylation patterns across individual chromatin fibers ^13^ (**Figure 1A**). This preserves information about accessibility, nucleosome protection, and footprints on the same molecule. We used this information to classify chromatin fibers according to their local methylation patterns rather than relying on population-averaged profiles. For each genomic feature class, molecules were aligned to a common reference point, and single-molecule GpC methylation profiles were used to capture local accessibility within a 2kb region, protected intervals, and nucleosome phasing. Single molecules were pooled across samples and clustered using the Leiden algorithm ^14^, producing common chromatin states that could be compared across samples. The resulting states were annotated using independent genomic datasets, including ChromHMM annotations and ChIP-seq enrichment for binding profiles of CTCF, SMC3 and TAD boundaries. This workflow allowed us to ask whether perturbation of cohesin redistributes chromatin fibers among shared local chromatin states.

**Figure 1.**
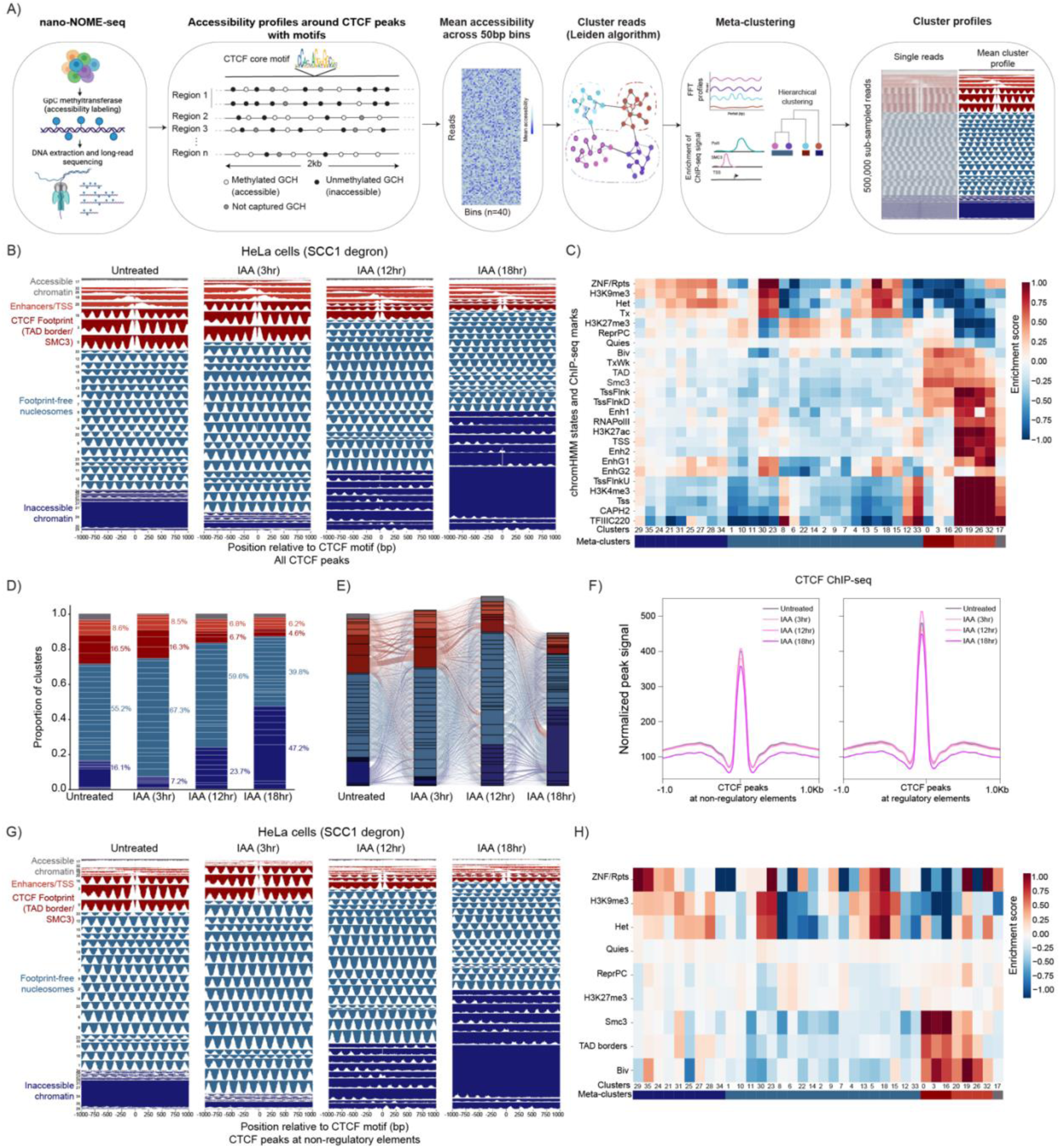
Acute SCC1/cohesin depletion leads to reorganization of chromatin states in HeLa cells. **A)** Overview of nano-NOME-seq analysis workflow to align and cluster reads to CTCF sites with motifs. **B)** Mean cluster profiles across time course depletion of SCC1 in HeLa cells. Clusters are colors based on meta-cluster assignment and annotation labels based on enrichment profiles. **C)** Heatmap of enrichment scores for each cluster based on chromHMM annotated states and ChIP-seq peaks. **D)** Barplot of the proportion of clusters across time points for SCC1 depletion. Percentage of meta-clusters is displayed. **E)** Alluvial plot showing the transition across the time points based on the most predominant meta-cluster at each CTCF site. **F)** CTCF ChIP-seq profiles (RPKM normalization) at CTCF peaks across the time points. CTCF peaks were separated based on overlap with and without regulatory elements. **G)** Mean cluster profiles based only on reads that align with CTCF sites that do not overlap with regulatory elements. **H)** Heatmap of enrichment score based on the subset of reads that align with CTCF sites that do not overlap with regulatory elements.

### SCC1 depletion progressively disrupts CTCF-centered chromatin states

We first applied this framework to CTCF-centered sites containing the core CTCF motif during acute SCC1/cohesin depletion. The nano-NOMe states captured a spectrum of CTCF-centered chromatin organization. Strong footprint states corresponding to dark-red clusters were enriched for CTCF, SMC3, TAD boundaries, and bivalent-chromatin annotations and showed phased flanking nucleosomes (**Figure 1B, C**). In contrast, weaker footprint states were more enriched for enhancer/TSS-associated annotations and showed reduced SMC3 enrichment (light red clusters), suggesting that CTCF sites embedded in active regulatory contexts adopt distinct configurations from boundary-like chromatin. In addition, we detected nucleosome phased states (light blue clusters) that lack a footprint and are enriched for different types of elements such as regulatory elements (clusters 12, 33) and polycomb/heterochromatin (the remaining light blue clusters). Finally, we also detected inaccessible molecules with low nucleosome occupancy (dark blue clusters) that are enriched for heterochromatin.

Across the SCC1-depletion time course, both regulatory-element-associated and boundary-associated CTCF states showed progressive loss of footprint integrity. Regulatory-element-associated states lost the already weaker CTCF-centered protection, while boundary-associated dark-red states showed erosion of the sharp footprint and reduced flanking nucleosome phasing (**Figure 1B**). Quantification of state proportions confirmed this redistribution, with SCC1 depletion reducing ordered CTCF/cohesin -associated states and increasing intermediate, disordered, inaccessible states over time (**Figure 1D**). The state transitions at individual CTCF sites based on the predominant chromatin state further highlights a progressive redistribution of molecules away from ordered CTCF-footprint states toward footprint-weakened, footprint-free, and inaccessible configurations across the depletion time course (**Figure 1E**). Thus, cohesin loss shifts individual CTCF-centered molecules away from ordered chromatin states. Importantly, this single-molecule reorganization occurred without a proportional loss of aggregate CTCF ChIP-seq signal, indicating that cohesin helps maintain local nucleosome order at CTCF-bound sites rather than simply preserving CTCF occupancy (**Figure 1F**).

Because CTCF sites overlapping promoters, enhancers, or other regulatory annotations may carry pre-existing regulatory chromatin architecture, we next repeated the analysis after excluding CTCF sites associated with regulatory elements. CTCF sites without regulatory elements were enriched for boundary-like CTCF elements and displayed a distinct CTCF-centered footprint architecture, characterized by sharp central protection and regular flanking nucleosome phasing (**Figure 1G, H**). Upon SCC1 depletion, this boundary-enriched architecture shifted toward footprint-weakened and less ordered states, indicating that cohesin-dependent disruption is not confined to promoter- or enhancer-associated CTCF sites. Notably, some light-red clusters persisted even after removal of annotated CTCF sites at regulatory elements. These clusters retained partial CTCF-centered protection but showed footprint patterns resembling those seen at TSS- or enhancer-associated CTCF sites, suggesting that unannotated or cell-type-specific cofactors may shape local chromatin architecture at a subset of CTCF sites, consistent with our previous finding that transcription-factor context modulates CTCF-associated nucleosome organization ^9^. These clusters showed reduced enrichment for SMC3 and TAD-boundary annotations, but increased enrichment for bivalent chromatin marks, compared to the dark red boundary CTCF clusters.

We next asked whether the stronger footprinting and nucleosome phasing at CTCF sites that do not overlap with regulatory elements could be explained by stronger CTCF binding. This was not the case: aggregate CTCF ChIP-seq profiles showed comparable or stronger CTCF signal at CTCF sites that overlap with regulatory elements than at sites not associated with regulatory elements across the SCC1-depletion time course (**Figure 1F**). Thus, the greater footprint strength and flanking nucleosome order at boundary-enriched CTCF sites cannot be attributed to higher bulk CTCF signal. This dissociation highlights an important point: the sites with the strongest boundary-like single-molecule nucleosome architecture are not necessarily those with the strongest aggregate CTCF occupancy. Conversely, CTCF sites embedded in regulatory chromatin can retain strong CTCF ChIP-seq signal while showing weaker footprinting and reduced nucleosome phasing. These data show that CTCF ChIP-seq peak strength is insufficient to predict local footprint integrity or nucleosome phasing, consistent with our previous finding that local chromatin context modulates CTCF-associated nucleosome organization ^9^.

Together, the data show that CTCF sites that do not overlap with regulatory elements define a boundary-enriched chromatin class whose ordered nucleosome architecture depends on cohesin. Although CTCF sites at regulatory elements showed comparable or stronger aggregate CTCF ChIP-seq signal, they displayed weaker footprinting and reduced nucleosome phasing, indicating that CTCF signal strength does not explain the boundary-like chromatin state. Thus, CTCF binding can persist in aggregate while SCC1 depletion progressively disrupts the surrounding nucleosome architecture, indicating that cohesin is required to maintain the ordered CTCF-centered boundary state.

### CTCF-centered chromatin states are redistributed across the cell cycle

Because cohesin residence time and function change across the cell cycle, we next asked whether CTCF-centered chromatin states vary between G1, S, G2, and cells in mitosis. During S phase, replication-coupled cohesin acetylation promotes Sororin association with cohesin, which antagonizes WAPL-mediated release and stabilizes a cohesive post-replicative pool; thus, S/G2 contains both dynamic extrusion-associated cohesin and Sororin-stabilized cohesive cohesin ^10^. Molecules from all cell-cycle conditions were again pooled before clustering, allowing the same CTCF-centered states to be compared across phases rather than redefining clusters independently in each population. This analysis recovered the same broad spectrum of local chromatin configurations described in **Figure 1** (**Figure 2A**).

**Figure 2.**
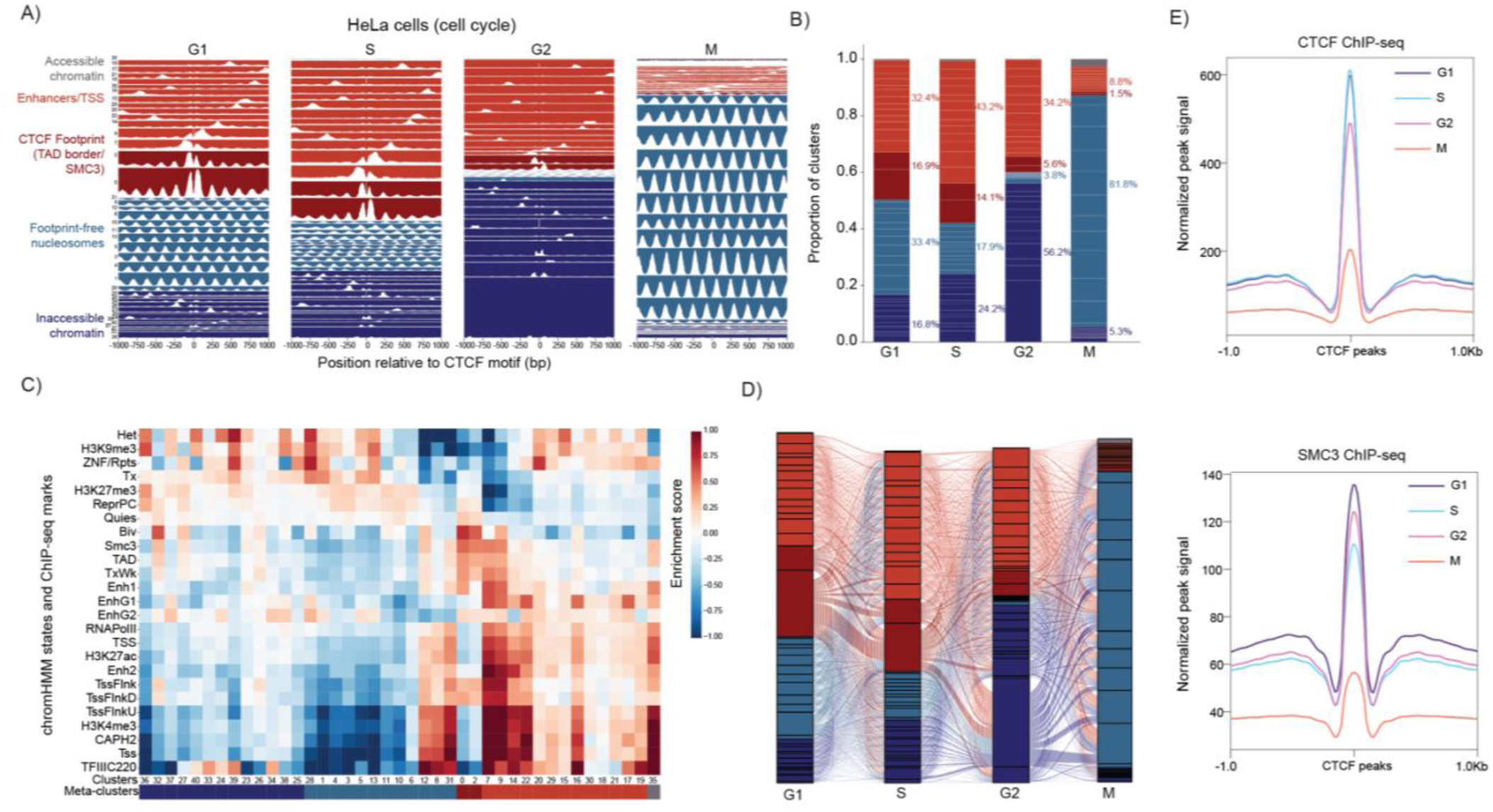
Reorganization of chromatin states across the cell cycle phases in HeLa cells. **A)** Mean cluster profiles across cell cycle phased in HeLa cells. **B)** Barplot of the proportion of clusters across cell cycle phased. Percentage of meta-clusters is displayed. **C)** Heatmap of enrichment scores for each cluster based on chromHMM annotated states and ChIP-seq peaks. **D)** Alluvial plot showing the transition across the cell cycle phased based on the most predominant meta-cluster at each CTCF site.

Cell-cycle progression caused a pronounced redistribution of molecules among CTCF-centered states. In G1, cells retained prominent CTCF-footprint states with phased flanking nucleosomes, consistent with ordered boundary organization during a phase dominated by extrusion-associated cohesin. In S and G2, when cohesin exists in both extrusive and cohesive pools, molecules shifted away from uniformly ordered boundary configurations toward more heterogeneous and inaccessible chromatin states, marked with a weak regulatory element signature (light red clusters) and inaccessible heterochromatin marks (dark blue clusters) (**Figure 2A-D**). We next compared the chromatin state changes with population-averaged CTCF and SMC3 ChIP-seq signal across the same cell-cycle stages (**Figure 2E**). Because aggregate CTCF and SMC3 occupancy are comparatively maintained during interphase, this redistribution is best interpreted as a change in the chromatin state surrounding CTCF-bound sites, rather than a simple consequence of factor loss. Mitotic cells were qualitatively distinct: most molecules occupied footprint-free nucleosomal states that were depleted of active marks and weakly enriched for heterochromatin and Polycomb features. Consistent with this mitotic state shift, ChIP-seq signal was reduced in mitosis. These cells showed little central CTCF protection, consistent with the reported loss of site-specific CTCF binding in prometaphase and the prophase-pathway removal of most cohesin from chromosome arms ^15–17^. Thus, the mitotic CTCF-centered state likely reflects both reduced CTCF/cohesin occupancy and replacement of interphase boundary organization by a nucleosome-dominated architecture.

Together, these data show that CTCF-centered chromatin organization is progressively remodeled across the cell cycle. As cells enter S and G2, CTCF-centered sites become less uniformly organized and more frequently adopt inaccessible chromatin configurations as cohesin is redistributed from a predominantly extrusive G1 pool toward mixed extrusive and cohesive states. By mitosis, most cohesin has been removed from chromosome arms through the prophase pathway, while the limited cohesin that remains on arms is thought to be extrusion-competent. This remodeling coincides with a footprint-free but phased nucleosome state, indicating that local CTCF-centered organization is not simply erased but replaced by a nucleosome-dominated architecture as CTCF binding is reduced and cohesin is largely depleted from chromosome arms.

These results indicate that ordered CTCF-centered chromatin is a cell-cycle-regulated state. The persistence of ordered footprint states in G1 supports the idea that local boundary organization is linked to extrusion-associated cohesin activity, whereas the altered state distribution in G2 suggests that the emergence of a Sororin-stabilized cohesive pool is not sufficient to preserve the same ordered CTCF-centered architecture. This distinction motivated us to test whether Sororin-stabilized cohesin in G2 contributes to, or instead separates from, the local chromatin states that organize CTCF sites.

### Sororin-stabilized cohesin is associated with reduced CTCF-centered footprinting in G2

To test whether the G2 chromatin-state redistribution reflects the increasing contribution of Sororin-stabilized cohesive cohesin, we acutely depleted Sororin in mESCs and compared the resulting G2-enriched population with RO-3306-treated G2 control cells. Control cells were collected after RO-3306 treatment to enrich for G2, whereas 10 h IAA-mediated Sororin depletion efficiently reduced Sororin-AID/eGFP and caused Sororin-degron cells to accumulate with a predominantly G2 DNA-content profile (**Figure 3A**). This allowed us to examine CTCF-centered chromatin organization after loss of Sororin-stabilized cohesive cohesin. We then applied the same CTCF-centered single-molecule clustering framework used above for HeLa cells. Because mESCs have a more open chromatin landscape than HeLa, they showed a different baseline distribution of single-molecule states, including greater representation of accessible regulatory-element-associated states and more CTCF-footprinted clusters overall. We therefore interpreted Sororin-dependent changes as redistribution within the mESC CTCF-centered state space, rather than as a direct one-to-one match to the HeLa cluster distribution.

**Figure 3.**
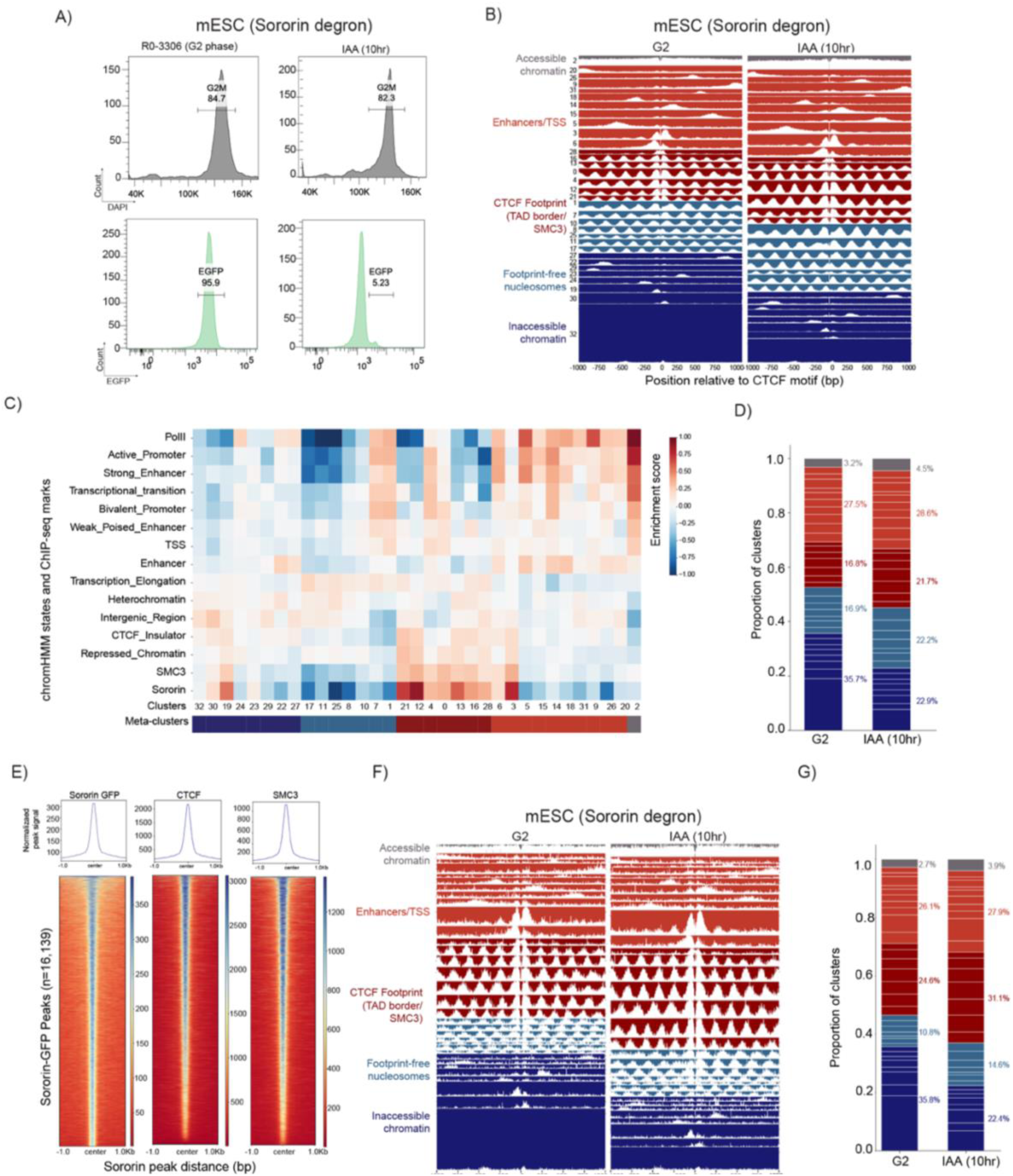
Depletion of sororin in mESCs leads to an increase in ordered foot-print chromatin state in G2 phase. **A)** Flow cytometry profiles of DAPI staining (DNA content, upper panels) and GFP fluorescence (Sororin expression, lower panels) in Sororin-AID mESCs following treatment with 500 μM IAA for 10 h or 10 μM RO-3306 for 14 h. **B)** Mean cluster profiles in G2 phase and sororin depleted cells. **C)** Heatmap of enrichment scores for each cluster based on chromHMM annotated states and ChIP-seq peaks. **D)** Barplot of the proportion of clusters in G2 and sororin depleted cells. Percentage of meta-clusters is displayed. **E)** ChIP-seq profiles for sororin (GFP-tagged), CTCF and SMC3 at Sororin-GFP peaks. **F)** Mean cluster profiles in G2 phase and sororin depleted cells for reads that align with CTCF peaks that overlap with sororin and SMC3. **G)** Barplot of the proportion of clusters in a subset of reads that align with CTCF peaks that overlap with sororin and SMC3. Percentage of meta-clusters is displayed.

Untreated G2 mESCs showed a relative reduction in ordered CTCF-centered architecture and greater representation of inaccessible or footprint-free nucleosomal states compared to G1, consistent with the G2 shift observed above. Sororin depletion reduced these inaccessible states and increased CTCF-footprinted clusters enriched for Sororin and SMC3 annotations, shifting G2 CTCF-centered molecules toward boundary-associated configurations (**Figure 3B-D**). Thus, Sororin depletion shifts G2 CTCF-centered molecules toward more footprinted boundary-like configurations, supporting a model in which CTCF-centered nucleosome order is favored by dynamic, extrusion-associated cohesin rather than Sororin-stabilized cohesive cohesion.

We next asked whether this effect was most evident at sites directly occupied by Sororin-associated cohesin. Sororin-GFP peaks were strongly enriched for CTCF and SMC3, confirming that Sororin is concentrated at a subset of CTCF/cohesin sites in G2 (**Figure 3E**). However, this relationship was highly asymmetric: Sororin marked only 11.2% of CTCF peaks, yet among SMC3-overlapping Sororin peaks, ∼95% overlapped CTCF and only ∼5% overlapped SMC3 without CTCF. This minor SMC3-only fraction may represent CTCF-independent, Sororin-stabilized cohesin at promoters, enhancers, or other noncanonical retention sites, although weak CTCF occupancy below detection cannot be excluded. Centering the single-molecule analysis on Sororin-GFP peaks resulted in a larger increase of the dark red CTCF-footprinted clusters in the sororin depleted cells (**Figure 3F, G**). Thus, the sites directly occupied by Sororin-associated cohesin are precisely those that become more CTCF-footprinted after Sororin loss. These results indicate that removal of Sororin shifts Sororin-bound chromatin toward a broader set of footprinted CTCF-centered states, consistent with extrusion-competent cohesin being more relevant for generating or maintaining these architectures.

Together, the cell-cycle and Sororin-depletion analyses separate cohesive cohesin from extrusion-associated cohesin at CTCF sites. Although mESCs show more CTCF-footprinted states overall than HeLa cells, the G2 population still shows a relative shift away from the ordered, footprinted architectures enriched in G1 and toward more inaccessible configurations. Depleting Sororin shifts this balance back toward CTCF-footprinted states. These findings support the conclusion that CTCF-centered chromatin order is linked to cohesin-barrier encounters during extrusion, whereas Sororin-stabilized cohesive cohesin is associated with reduced representation of these footprinted architectures in G2.

### TSS-centered chromatin states are remodeled across the cell cycle and after cohesin depletion

We next asked whether cohesin-dependent chromatin-state remodeling extends beyond CTCF-centered boundary elements to transcription start sites. TSS-centered molecules were pooled across cell-cycle phases and clustered using the same single-molecule nano-NOMe-seq framework, recovering accessible promoter/TSS states, CTCF-footprinted SMC3/boundary-associated states, footprint-free phased nucleosomal states, and inaccessible chromatin states (**Figure 4A-C**).

**Figure 4.**
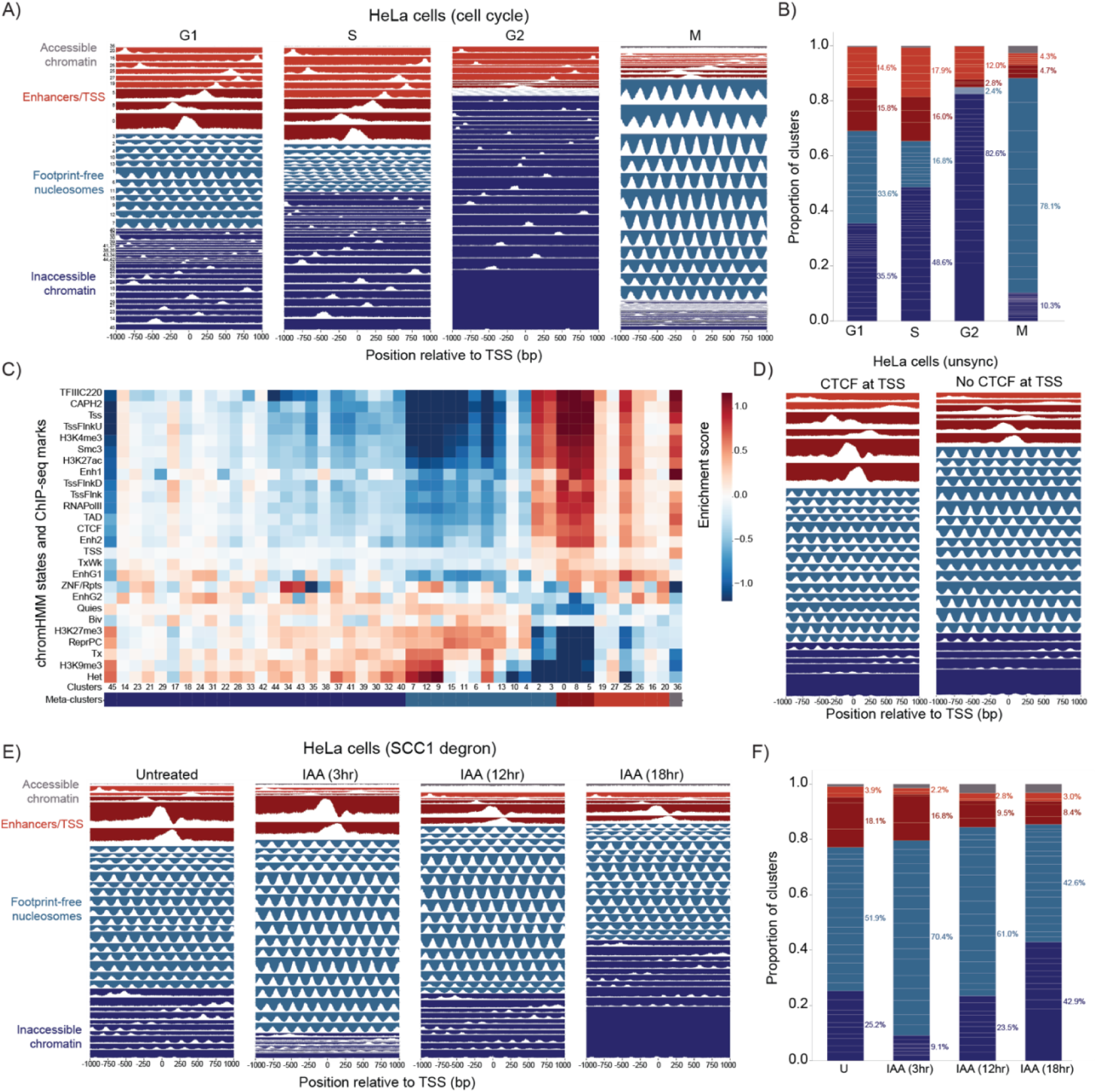
Chromatin states at transcription start sites across cell cycle phases and SCC1 degradation time points in HeLa cells. **A)** Mean cluster profiles across cell cycle phased in HeLa cells based on read clustering at transcription start sites. **B)** Barplot of the proportion of clusters across cell cycle phased. Percentage of meta-clusters is displayed. **C)** Heatmap of enrichment scores for each cluster based on chromHMM annotated states and ChIP-seq peaks. D) Mean cluster profiles separated based on reads that align with TSSs with CTCF and TSSs without CTCF. **E)** Mean cluster profiles across time points for SCC1 degradation in HeLa cells based on read clustering at transcription start sites. **F)** Barplot of the proportion of clusters across time points. Percentage of meta-clusters is displayed.

In G1, these states were broadly represented across promoters. As cells progressed through S and G2, the distribution shifted toward increasingly inaccessible chromatin states, indicating that promoter-centered chromatin organization is also cell-cycle regulated. Mitotic TSSs were qualitatively distinct, with most molecules occupying footprint-free phased nucleosomal states and showing reduced representation of accessible or footprinted promoter configurations (**Figure 4A, B**). Thus, as at CTCF-centered sites, mitosis replaces interphase promoter states with a nucleosome-dominated architecture rather than simply eliminating local organization.

We then stratified promoters by the presence or absence of promoter-proximal CTCF. This revealed distinct chromatin-state distributions at CTCF-positive and CTCF-negative TSSs. CTCF-positive TSSs were enriched for accessible enhancer/TSS-associated states and footprint configurations, whereas CTCF-negative TSSs were dominated by footprint-free phased nucleosomal states (**Figure 4D**). Thus, promoter-proximal CTCF changes the balance of TSS-centered chromatin states but does not simply impose a canonical boundary-like architecture. Instead, CTCF-positive promoters occupy a more accessible regulatory chromatin environment, while CTCF-negative promoters retain ordered nucleosome arrays without a promoter-proximal CTCF footprint.

Finally, we tested whether cohesin contributes to these promoter-associated states by analyzing TSS-centered molecules during SCC1 depletion. Acute SCC1 loss progressively remodeled promoter chromatin (**Figure 4E, F**). Early depletion shifted molecules toward footprint-free phased nucleosomal states, whereas prolonged depletion reduced regulatory-associated/footprinted states and increased inaccessible chromatin configurations. Within the dark-red regulatory-associated clusters, we also observed footprinted promoter architectures that may reflect distinct Pol II-associated states, including configurations with a promoter-proximal footprint and positioned +1 nucleosome consistent with pausing, and others with more downstream protection patterns consistent with early elongation. These putative paused- and elongation-associated configurations remained proportionally represented after SCC1 depletion, indicating that SCC1 loss does not selectively collapse one of these promoter-footprinted states. Instead, cohesin depletion primarily altered the broader balance between regulatory-associated, footprint-free nucleosomal, and inaccessible promoter configurations. These dynamics indicate that cohesin helps maintain the normal distribution of promoter-centered chromatin states.

Together, these analyses show that cohesin-dependent single-molecule chromatin organization is not restricted to CTCF boundary elements. Promoters also exist in distinct chromatin states whose distribution depends on cell-cycle phase, promoter-proximal CTCF, and SCC1/cohesin. Thus, cohesin contributes to local nucleosome organization at both architectural boundaries and transcription start sites, while promoter-proximal CTCF defines a more accessible enhancer/TSS-associated promoter state rather than simply converting promoters into canonical boundary elements.

## Discussion

Cohesin is widely understood as the motor that builds loops and contact domains, but our results show that cohesin also maintains local nucleosome architecture at the sites where extrusion is positioned or constrained, including CTCF boundary elements and transcription start sites. Using single-molecule nano-NOMe-seq, we find that CTCF sites and TSSs do not adopt uniform chromatin architectures, but instead comprise distinct nucleosome configurations that differ in CTCF footprint strength, nucleosome phasing, accessibility, regulatory annotation, and cohesin association. Acute SCC1 depletion progressively shifts molecules away from ordered CTCF-footprinted states toward footprint-weakened, footprint-free, and inaccessible configurations. Thus, cohesin is required not only for long-range genome folding, but also for maintaining the local nucleosome architecture surrounding CTCF-bound boundary sites and promoters.

A central conclusion from this work is that CTCF occupancy alone does not define an ordered boundary state. CTCF sites away from regulatory elements showed the strongest CTCF-centered footprints and most regular flanking nucleosome phasing, consistent with a boundary-enriched chromatin class. However, this stronger local organization was not explained by higher aggregate CTCF ChIP-seq signal. Indeed, CTCF sites at regulatory elements showed stronger CTCF signal despite weaker footprinting and reduced nucleosome phasing. This distinction supports a model in which CTCF-bound sites adopt different single-molecule chromatin states depending on local regulatory context and cohesin engagement. In this view, boundary strength is not encoded simply by CTCF binding intensity, but by the fraction of molecules at a site that occupy an ordered, cohesin-dependent CTCF-centered nucleosome architecture.

These findings also suggest why architectural perturbations can have uneven regulatory consequences. Contact maps and ChIP-seq profiles average across many molecules, whereas boundary and promoter function may depend on the fraction of molecules occupying particular local chromatin states. A CTCF site can retain aggregate binding while losing the phased nucleosome architecture associated with strong boundaries, and promoters with or without nearby CTCF can differ in their balance of footprinted, accessible, and phased nucleosomal configurations. Thus, single-molecule chromatin-state composition provides information that is not captured by CTCF peak height, average accessibility, or contact frequency alone.

The cell-cycle analysis further indicates that CTCF-centered chromatin organization is dynamically regulated. G1 cells retain prominent ordered CTCF-footprint states with phased flanking nucleosomes, consistent with a phase in which cohesin is largely available for extrusion-associated boundary encounters. As cells progress through S and G2, CTCF-centered sites shift toward more heterogeneous and inaccessible configurations as cohesin is redistributed from a predominantly extrusive G1 pool toward mixed extrusive and cohesive states. However, the magnitude and composition of this shift depend on the baseline chromatin environment of each cell type. mESCs, which have a more open chromatin landscape, retain a higher proportion of CTCF-footprinted molecules than HeLa cells, whereas HeLa cells show a greater representation of inaccessible or disordered configurations. Thus, cell-cycle progression drives a common shift away from ordered CTCF-centered architecture, but the final state distribution is shaped by cell-type-specific chromatin accessibility and starting nucleosome organization. By mitosis, most cohesin has been removed from chromosome arms through the prophase pathway, while the limited remaining arm-associated cohesin is thought to be extrusion-competent. In this context, CTCF-centered sites lose central protection as CTCF occupancy is reduced and most cohesin is removed from chromosome arms. The resulting mitotic state is therefore not an intact boundary state, but a footprint-free phased nucleosomal architecture that replaces interphase CTCF/cohesin-centered organization.

The Sororin experiments separate cohesive cohesin from the cohesin activity most closely associated with CTCF-centered chromatin order. Sororin stabilizes post-replicative cohesive cohesin in S/G2, and Sororin depletion shifted G2 CTCF-centered molecules toward more footprinted boundary-like configurations. Specifically, loss of Sororin reduced inaccessible configurations and increased CTCF-footprinted molecules, including SMC3/TAD-boundary-enriched clusters and additional footprinted states. This redistribution suggests that ordered CTCF-centered nucleosome architecture is favored when cohesin is available for dynamic boundary encounters rather than stabilized primarily in a cohesive G2 pool. This effect was also observed when the analysis was centered specifically on Sororin-GFP peaks, indicating that Sororin-bound sites are the sites most affected by Sororin loss. These results suggest that Sororin-stabilized cohesive cohesin is associated with reduced representation of footprinted CTCF-centered states in G2, whereas extrusion-competent cohesin is more closely linked to ordered boundary-like configurations.

Together, the HeLa cell-cycle analysis and mESC Sororin experiments indicate that CTCF-centered nucleosome organization is dynamically regulated, but that the final distribution of molecules depends on cell-type-specific chromatin context. Dynamic, extrusion-associated cohesin is linked to ordered CTCF footprints and phased flanking nucleosomes, whereas accumulation of Sororin-stabilized cohesive cohesin in G2 is associated with a shift toward less ordered and more inaccessible CTCF-centered configurations. This shift is superimposed on the baseline chromatin environment of each cell type: mESCs retain more open chromatin and a higher proportion of CTCF-footprinted molecules than HeLa cells, but nevertheless show a G2-associated movement away from the ordered architectures enriched in G1. Thus, cell-cycle regulation of cohesin produces a common directional shift in CTCF-centered nucleosome organization, while the final distribution of molecules is shaped by cell-type-specific chromatin accessibility and the starting level of boundary-associated footprinting.

Our TSS-centered analysis extends this principle beyond canonical CTCF boundary elements. Promoters also displayed distinct single-molecule nucleosome architectures: some molecules were accessible and enhancer/TSS-associated, some carried CTCF-centered footprints, others formed footprint-free phased nucleosomal arrays, and others were largely inaccessible. These promoter configurations are redistributed across the cell cycle and after SCC1 depletion, indicating that promoter chromatin organization is also cohesin-sensitive. Stratifying TSSs by promoter-proximal CTCF revealed that CTCF-positive promoters are enriched for accessible enhancer/TSS-associated states, including regulatory-element-associated footprinted configurations, whereas CTCF-negative promoters are dominated by footprint-free phased nucleosomal states. Thus, promoter-proximal CTCF changes the balance of promoter chromatin states but does not convert promoters into canonical boundary elements.

The SCC1-depletion time course at TSSs suggests that cohesin contributes to the normal balance of promoter-centered architectures. Early SCC1 depletion shifted molecules toward footprint-free phased nucleosomal states, whereas prolonged depletion increased inaccessible chromatin configurations and reduced regulatory-associated states. Within the dark-red regulatory-associated clusters, we observed distinct footprinted promoter architectures that may correspond to different Pol II-associated configurations, including putative paused states with a positioned +1 nucleosome and more downstream protection patterns that could reflect early elongation. The presence of these configurations highlights the potential of single-molecule chromatin profiling to resolve promoter states that are averaged together in bulk assays. Importantly, these putative paused- and elongation-associated configurations remained proportionally represented after SCC1 depletion, suggesting that cohesin loss primarily alters the broader distribution of promoter nucleosome configurations rather than selectively eliminating one promoter-footprinted substate.

Together, these findings support a model in which cohesin collisions act as local chromatin-patterning events. At boundary-enriched CTCF sites, extrusion-associated cohesin encounters CTCF and helps maintain a sharply footprinted, phased nucleosome architecture. At regulatory-element-associated CTCF sites and promoters, cohesin encounters occur in a different chromatin environment, producing more heterogeneous accessible and footprinted states. When SCC1 is depleted, these ordered local architectures progressively erode even when aggregate CTCF signal persists. As cohesin becomes increasingly Sororin-stabilized in G2, CTCF-centered footprint states are reduced, whereas Sororin depletion shifts molecules back toward more footprinted configurations. Thus, cohesin-dependent local organization reflects the functional state and positioning of cohesin, not cohesin occupancy alone.

Overall, our results identify local nucleosome organization as a cohesin-maintained feature of both CTCF boundaries and promoters. They suggest that the functional output of cohesin is not limited to the formation of loops and domains but includes the maintenance of ordered chromatin states at sites where extrusion is halted, slowed, or positioned. This provides a mechanistic framework for understanding why CTCF occupancy, cohesin occupancy, and transcriptional output are often imperfectly coupled: the critical regulatory variable may be the fraction of molecules that retain an ordered cohesin-dependent local architecture. In this model, cohesin collisions pattern chromatin locally, linking the mechanics of loop extrusion to the single-molecule chromatin states that underlie boundary function and promoter organization.

## Acknowledgements

This work was supported by 1R35GM122515 (J.S) and NIH P01CA229086 (J.S). GC and GJ were supported by fellowships from the NCC.

The authors thank Skok lab members for helpful scientific discussions, New York University School of Medicine High Performance Computing Facility (HPCF) for computing technical support, Adriana Heguy and the Genome Technology Center (GTC) core for sequencing efforts, NYU Flow Cytometry and Cell Sorting Center for FACS analysis and sorting. GTC is a shared resource partially supported by the Cancer Center Support Grant P30CA016087 at the Laura and Isaac Perlmutter Cancer Center.

## Author contributions

These studies were designed by Jane Skok. The analysis was performed by Ramya Raviram and Tom Schippke. Experiments were performed by Guimei Jiang and Giulia Cova. The paper was written by Jane Skok and Ramya Raviram.

## METHODS

### Cell lines and culture conditions

HeLa S3 cells were derived from the line described previously by Abramo *et al.* ^18^. HeLa SCC1-AID cells were from Wutz *et al.* ^19^. SORORIN-AID-eGFP; Tir1(Rosa26); RAD21-HaloTag mESCs were described by Nora *et al.* ^20^.

HeLa S3 and HeLa SCC1-AID cells were cultured in DMEM (Gibco, 11965-092) supplemented with 10% DBS (Gibco, 10371-029), 1% L-glutamine (Gibco, 25030-081), 1% penicillin–streptomycin (Gibco, 15070-063) and 1% MEM non-essential amino acids (Gibco, 11140-050). For auxin-induced degradation, HeLa SCC1-AID cells were treated with 500 μM indole-3-acetic acid sodium salt (IAA; Sigma, I5148) for the indicated times: 0, 3, 12 and 18 h.

Mouse E14Tg2a embryonic stem cells (karyotype 19, XY; 129/Ola isogenic background) were cultured feeder-free on 0.1% gelatin-coated dishes (Sigma, ES-006-B; Falcon, 353003) at 37 °C and 5% CO₂ in a humidified incubator. Cells were maintained in DMEM (Thermo Fisher, 11965-118) supplemented with 15% fetal bovine serum (Thermo Fisher, SH30071.03), 100 U ml⁻¹ penicillin and 100 μg ml⁻¹ streptomycin (Sigma, P4458), 1× GlutaMAX (Thermo Fisher, 35050-061), 1 mM sodium pyruvate (Thermo Fisher, 11360-070), 1× MEM non-essential amino acids (Thermo Fisher, 11140-050), 50 μM β-mercaptoethanol (Sigma, 38171), 10⁴ U ml⁻¹ leukemia inhibitory factor (Millipore, ESG1107), 3 μM CHIR99021 (Sigma, SML1046) and 1 μM MEK inhibitor PD0325901 (Sigma, PZ0162). Cells were passaged every other day using TrypLE (Thermo Fisher, 12563011).

For Sororin depletion, SORORIN-AID-eGFP; Tir1(Rosa26); RAD21-HaloTag mESCs were treated with 500 μM IAA (Sigma, I5148-2G) for 10 h, using a 1,000× filter-sterilized stock prepared in water.

### HeLa cell-cycle synchronization

For G1 synchronization, 1.5 × 10⁶ HeLa S3 cells were seeded overnight in 10-cm dishes. Cells were treated with 2 mM thymidine for 16 h, washed with PBS, released into thymidine-free medium for 8 h, subjected to a second 2 mM thymidine block for 16 h and then released into thymidine-free medium for 13 h before collection.

For S-phase synchronization, 1.5 × 10⁶ HeLa S3 cells were seeded overnight in 10-cm dishes. Cells were treated with 2 mM thymidine for 16 h, washed with PBS, released into thymidine-free complete medium for 8 h, subjected to a second 2 mM thymidine block for 16 h and then released into thymidine-free medium for 2 h before collection.

For G2 synchronization, 1.5 × 10⁶ HeLa S3 cells were seeded overnight in 10-cm dishes. Cells were treated with 2 mM thymidine for 16 h, washed with PBS, released into thymidine-free complete medium for 8 h, subjected to a second 2 mM thymidine block for 16 h and then released into thymidine-free medium for 6 h before collection.

For mitotic synchronization, HeLa S3 cells were synchronized by double thymidine block followed by nocodazole treatment and mitotic shake-off. Cells were treated with 2 mM thymidine for 16 h, washed with PBS, released into thymidine-free medium for 8 h, subjected to a second 2 mM thymidine block for 16 h and then released into thymidine-free medium for 8 h. Cells were then treated with 300 nM nocodazole for 2 h, and mitotic cells were collected by shake-off.

HeLa S3 cells were not sorted after synchronization at G1, S, G2 or M phase. At collection, an aliquot of each synchronized culture was fixed with 4% paraformaldehyde and stained with DAPI to assess synchronization efficiency. Only samples passing this quality-control assessment were used for downstream experiments.

### mESC synchronization

For G2 phase synchronization SORORIN-AID-eGFP mES cells were seeded for 10–12 h and then treated with 10 μM RO-3306 for 14 h. Cells were collected at the end of treatment for downstream analyses. All incubations were performed in complete medium at 37 °C in a humidified incubator with 5% CO₂ unless otherwise stated.

### Nano-NOMe-seq

1 × 10^6 cells were collected, washed in cold PBS, and nuclei were isolated as described in Battaglia et al., ^21^ using cold lysis buffer (10 mM Tris-HCl pH 7.4, 10 mM NaCl, 0.5 mM spermidine, 1.5 mM MgCl2, 0.1 mM EDTA, 0.25% IGEPAL CA-630) for 3–6 min, followed by washing in nuclei wash buffer (10 mM Tris-HCl pH 7.4, 50 mM NaCl, 0.5 mM spermidine, 1.5 mM MgCl2, 0.1 mM EDTA).

For GpC methyltransferase labeling, nuclei were resuspended in 1× GpC MTase reaction buffer and equilibrated to 37°C. Methylation was performed in two sequential 6-min incubations with M.CviPI and SAM. Reactions were stopped with prewarmed 2× stop buffer (20 mM Tris-HCl pH 7.4, 600 mM NaCl, 10 mM EDTA, 1% SDS), followed by proteinase K digestion (200 μg/mL, 4 h at 56°C), RNase A treatment (100 μg/mL, 30 min at 37°C), and a second proteinase K digestion (200 μg/mL, 30 min at 37°C).

High-molecular-weight DNA was extracted by phenol-chloroform, precipitated with 1 volume of isopropanol, washed in 70% ethanol, resuspended in EB buffer, and quantified by Nanodrop 2000. DNA size was assessed using Agilent genomic DNA ScreenTape.

### Nano-NOMe-seq library preparation

For Nano-NOMe-seq, DNA was sheared to ∼20 kb using a Covaris G-Tube, size-checked by Agilent ScreenTape, and 1.5 μg DNA was used for library preparation with the Oxford Nanopore SQK-LSK114 kit. Libraries were quantified by Qubit dsDNA HS, loaded at 150–200 fmol onto PromethION R10.4.1 flow cells, and sequenced on a PromethION P24 for 72 h. Runs used standard settings with rejection of fragments <1 kb, detection of 5mC and 5hmC, and super-accurate basecalling enabled.

### Nano-NOME-seq analysis

#### Pre-processing

Basecalling was performed from pod5 file using dorado ^22^ (v1.2.0) for the sup, 5mc and 5hmc modifications and --min-qscore of 10. We then aligned the reads using dorado aligner to the mm10 or hg38 genome. Aligned reads were then filtered for a mapq score of 10 and sorted using samtools (v1.2). Methylation calls were extracted using modkit ^23^ (version 0.5.0) for GCH sites in the hg38 or mm10 genome.

#### Alignning reads at CTCF and TSS sites

Reads for clustering analysis were filtered based on the following criteria 1) overlap with CTCF sites that contain the core motif (+/-1kb) or TSS (+/- 1kb), 2) have at least 100 GCH captured sites (methylated or unmethylated) with 80% base call probability. Annotation for TSS was based on the GENCODE annotation for mm10 (M25) and hg38 (v49). Reads mapping to genes transcribed in antisense direction were realigned to the sense direction. Similarly, for CTCF motifs detected on the antisense strand, reads were realigned to the sense direction.

#### Clustering reads

GCH accessibility signal for each read was binned into 50 bp windows across the 2 kb region and summarized using the mean methylation value per bin. Remaining missing values after binning were imputed using K-nearest neighbor imputation (k = 5, scikit-learn). An equal number of reads was sampled from each condition to ensure balanced representation across samples. A K-nearest neighbor graph (k = 50, scikit-learn) was constructed on the binned accessibility matrix using the Manhattan distance metric. Edge weights were computed as affinities by applying an exponential kernel to the pairwise distances. Community detection was then performed on this graph using the Leiden algorithm (leidenalg) with the RBConfigurationVertexPartition function and a specified resolution parameter (HeLa Cell Cycle at CTCF: 2.1, HeLa Cell Cycle at TSS: 2.1, HeLa SSC1 at CTCF: 3, HeLa SSC1 at TSS: 3, mESC Sororin at CTCF: 2.2).

#### Enrichment heatmaps for chromHMM and ChIP-seq peaks

Enrichment of chromatin states and ChIP-seq peaks for each cluster was computed using a log2 observed-over-expected ratio. We used bedtools intersect to overlap reads with the annotation features. For each cluster–annotation pair, the observed proportion was defined as the total overlap (in base pairs) between reads of that cluster and a given annotation state, divided by the total length of reads in that cluster. The expected proportion was defined as the total overlap across all reads with that state, divided by the total length of all reads. Enrichment was then calculated as log2(observed proportion / expected proportion). For visualization, annotation states (rows) were ordered by hierarchical clustering (Euclidean distance, Ward’s linkage, scipy) and clusters (columns) were ordered based on method described below.

#### Re-ordering clusters for visualization

The previous clusters were then grouped into macro-clusters. In the first stage, the mean accessibility profile of each cluster was computed, and its Fast Fourier Transform (FFT) magnitude spectrum was extracted, retaining the first 10 mid-frequency components and skipping the DC and the two lowest-frequency components corresponding to broad-scale trends. These FFT features were then standardized and concatenated with selected enrichment features (CTCF, SMC3, PolII, TSS). Hierarchical clustering (Euclidean distance and Ward linkage, scipy) was then applied to the combined features to determine the order of clusters and macro-clusters were determined based on distance cut-off (HeLa Cell Cycle at CTCF: 5, HeLa Cell Cycle at TSS: 5, HeLa SSC1 at CTCF: 4, HeLa SSC1 at TSS: 3.6, mESC Sororin at CTCF: 3).

### ChIPmentation

After cell treatment and cell trypsinization, collected cells were divided in 10 million aliquots and resuspended in fresh 1X PBS (1million cells/1mL). For double cross linking 25mM EGS (ethylene glycol bis(succinimidyl succinate); Thermofisher #21565) were added and cells were put in rotation for 30 min at room temperature, followed by addition of 1% formaldehyde (Tousimis #1008A) for 10 min also in rotation at room temperature. Quenching was performed by adding glycine to a final concentration of 0.125 M followed by incubations of 5 min at room temperature in rotation. Fixed cells were washed twice with 5 mL of 1 X PBS containing 0.5% BSA and centrifuged at 3000 rpm for 5 min at 4°C. Pellets were finally resuspended in 500 μL 1X PBS containing 0.5% BSA, transferred to 1.5 mL Eppendorf and centrifuged at 3000rpm for 3 min at 4°C. Supernatant was completely removed, pellets were snap-frozen in liquid nitrogen and stored at −80°C. Fixed cells (10 million) were thawed on ice, resuspended in 350 μL ice-cold lysis buffer (10 mM Tris-HCl (pH 8.0), 100 mM NaCl, 1 mM EDTA (pH 8.0), 0.5 mM EGTA (pH 8.0), 0.1% sodium deoxycholate, 0.5% N-lauroysarcosine and 1X protease inhibitors) and lysed for 10 min by rotating at 4°C. Chromatin was sheared using a bioruptor (Diagenode) for 15 min (30 s on, 30 s off, high output level). 100 μL of cold lysis buffer and 50 μL of 10% Triton X-100 (final concentration of 1) were then added and the samples were centrifuged for 5 min at full speed at 4°C. Supernatant was collected, transferred to a new tube (Protein Low Binding tube) and shearing was continued for another 10 min, then the chromatin was quantified. SMC3 (abcam, ab9263), CTCF (cell Signaling Technology, 3418), GFP (Abcam, ab6556) and Rabbit IgG (abcam, ab37415) antibodies were bound to protein A magnetic beads by incubation on a rotator for 1 h at room temperature. 10 μL each of antibody was bound to 50 μL of protein-A magnetic beads (Dynabeads) and added to the sonicated chromatin for immunoprecipitation at 4°C overnight. Next day, samples were washed and tagmentation were performed as per the original ChIPmentation protocol. In short, the beads were washed successively twice in 500 μL cold low-salt wash buffer (20 mM Tris-HCl (pH 7.5), 150 mM NaCl, 2 mM EDTA (pH 8.0), 0.1% SDS, 1% Triton X-100), twice in 500 μL cold LiCl-containing wash buffer (10 mM Tris-HCl (pH 8.0), 250 mM LiCl, 1 mM EDTA (pH 8.0), 1% Triton X-100, 0.7% sodium deoxycholate) and twice in 500 μL cold 10 mM cold Tris-Cl (pH 8.0) to remove detergent, salts and EDTA. Then the beads were resuspended in 25 μL of the freshly prepared tagmentation reaction buffer (10 mM Tris-HCl (pH 8.0), 5 mM MgCl2, 10% dimethylformamide) and 1 μL Tagment DNA Enzyme from the Tagment DNA Enzyme and Buffer Kit (Illumina #20034198) and incubated at 37°C for 10 min in a thermomixer. Following tagmentation, the beads were washed successively twice in 500 μL cold low-salt wash buffer (20 mM Tris-HCl (pH 7.5), 150 mM NaCl, 2 mM EDTA (pH8.0), 0.1% SDS, 1% Triton X-100) and twice in 500 μL cold Tris-EDTA-Tween buffer (0.2% tween, 10 mM Tris-HCl (pH 8.0), 1 mM EDTA (pH 8.0)). Chromatin was eluted and de-crosslinked by adding 70 μL of freshly prepared elution buffer (0.5% SDS, 300 mM NaCl, 5 mM EDTA (pH 8.0), 10 mM Tris-HCl (pH 8.0) and 10 μg/ml proteinase K for 1 h at 55°C 850rpm and overnight at 65°C 850rpm. Next day, the supernatant was collected, transferred to new DNA Low Binding tubes and supplemented with an additional 30 μL of elution buffer. DNA was purified using MinElute Reaction Cleanup Kit (Qiagen #28204) and eluted in 20 μl. Purified DNA (20 μL) was amplified as per the ChIPmentation protocol (Schmidl, Rendeiro et al. 2015) using indexed and non-indexed primers and NEBNext High-Fidelity 2X PCR Master Mix (NEB M0541) in a thermomixer with the following program: 72°C for 5 m; 98°C for 30 s; 14 cycles of 98°C for 10 s, 63°C for 30 s, 72°C for 30 s and a final elongation at 72°C for 1 m. DNA was purified using sparQ PureMag Beads (Quantabio, 95196 -060) to remove fragments larger than 700 bp as well as the primer dimers. Library quality and quantity were estimatred using Tapestation (Agilent High Sensitivity D1000 ScreenTape #5067–5584 and High Sensitivity D1000 reagents #5067–5585) and quantified by Qubit (Life Technologies Qubit 1X dsDNA High Sensitivity (HS) #Q33230). Libraries were then sequenced with the NovaSeq X+ Illumina technology according to the standard protocols and with around 200 million 150bp paired-end total per sample.

### ChIPmentation analysis

#### Pre-processing and peak calling

Paired-end reads were aligned to the hg38 or mm10 genome using bowtie2 (version 2.5.3) with the following arguments: --local --minins 25 --maxins 1500 --dovetail --no-discordant --no-mixed. Aligned reads were filtered for a mapq score of 30 using samtools (version 1.9) and PCR duplicates were filtered using sambamba (version 1.0.1). Peak calling was performed using macs2 using the narrow peak option for CTCF ChIP-seq data. bigWig files were created using the bamCoverage function from deeptools (version 3.5.5) and the RPKM normalization option.

#### CTCF motif overlap

CTCF peaks were filtered based on a p-value less than 1e-5 and the CTCF core motif was detected within peaks using fimo function from the MEME suit (version 5.5.7) using the MA0139.2 CTCF core motif. The default p-value cut-off of 1e-4 from fimo was used for detecting motif occurrence within a CTCF ChIP-seq peak.

